# Niche-aware metagenomic screening for enzyme methioninase illuminates its contribution to metabolic syntrophy

**DOI:** 10.1101/2024.05.03.592401

**Authors:** Erfan Khamespanah, Sedigheh Asad, Zeynab Vanak, Maliheh Mehrshad

## Abstract

The single step methioninase-mediated degradation of methionine (as a sulfur containing amino acid) is a reaction at the interface of carbon, nitrogen, sulfur, and methane metabolism in microbes. This enzyme is also a therapeutic target for its role in starving auxotrophic cancer cells. Applying our refined *in-silico* screening pipeline on 33,469 publicly available genome assemblies and 1878 MAGs/SAGs from brackish waters of the Caspian Sea and the Fennoscandian shield deep groundwater resulted in recovering 1845 methioninases. The majority of recovered methioninases belong to representatives of phyla Proteobacteria (50%), Firmicutes (29%), and Firmicutes_A (13%). Prevalence of methioninase among anaerobic microbes and in the anoxic deep groundwater together with the relevance of its products for energy conservation in anaerobic metabolism highlights such environments as desirable targets for screening novel methioninases and resolving its contribution to microbial metabolism and interactions. Among archaea, majority of detected methioninsaes are from representatives of *Methanosarcina* that are able to use methanethiol, the sulfur containing product from methionine degradation, as a precursor for methanogenesis. Branching just outside these archaeal methioninases in the phylogenetic tree we recovered 3 methioninases belonging to representatives of Patescibacteria reconstructed from deep groundwater metagenomes. We hypothesize that methioninase in Patescibacteria could contribute to their syntrophic interactions where their methanogenic partners/hosts benefit from the produced 2-oxobutyrate and methanethiol.

## Introduction

One commonly used and non-invasive approach to minimize the progression of cancer cells is to target their nutritional dependencies [1]. Since cancer cells have altered metabolism, they are not able to produce certain metabolites essential for their growth, thus depend on external supplies for such resources [2]. This phenomenon is specifically reminiscent of amino acid auxotrophies and different syntrophies detected in certain bacteria and archaea [3], [4], [5]. Amino acid auxotrophies in bacteria and archaea are hypothesized to be an evolutionary adaptation strategy to minimize the biosynthetic burden of carrying the pathway on the organism while simultaneously promoting cooperation between different taxa within the microbial community [6].

Depriving cancer cells of these essential biomolecules has been shown to hamper or in some cases block their growth, thus has become a desirable therapeutic approach in attenuating their proliferation [7]. It has been observed that most cancer cells depend upon external sources of different amino acids to support their increased rate of proliferation [8], [9], [10]. An example of targeting amino acid auxotrophies to manage cancer cells is the case of enzyme L-Asparaginase that has been used against Acute Lymphoblastic Leukemia (ALL), Since 1978, for its ability to indirectly deplete blood serum from accessible glutamine [11], [12], [13]. Another example would be arginine deaminase (ADI) that converts arginine to citrulline and ammonia and by doing so depletes this vital amino acid from the extracellular matrix of cancer cells [14], [15], [16]. More Recently, Methionine gamma-lyase (i. e., Methioninase), that breaks down methionine to 2-oxobutyrate, methanethiol, and ammonia, has become a therapeutic target due to its anti-tumor activity in cells that have become auxotrophic for methionine due to dysregulation of different pathways (**Supplementary Figure S1**) [17], [18], [19]. The Methioninase of the *Pseudomonas putida* that has been overexpressed as a recombinant protein in *Escherichia coli* is currently undergoing the first phase of clinical trial [20], [21].

Bacteria and archaea have been shown to be a promising resource to screen for amino acid degrading enzymes [22]. Optimum functionality of enzymes within their specific environmental conditions is the major contributor to the fitness of microorganisms in their respective niches [23]. The single step utilization of methionine, a sulfur containing amino acid, via the activity of methioninase [24], puts this reaction at the interface of carbon, nitrogen, sulfur, and methane metabolisms by producing end-products that could enter these metabolisms at different stages in microbes (**Figure 1**). This potentially hints at the specific niche of those microbial lineages carrying/expressing this enzyme. However, the eco-evolutionary significance of this enzyme, its range of functions, and distribution across different bacterial and archaeal lineages remain so far unexplored. Thus, target environments for screening this enzyme are not fully defined. Furthermore, since this enzyme is valuable for cancer treatment; it is expected that screening brackish environments that have salinity and osmolality similar to that of human serum; could potentially increases the possibility of finding enzymes compatible with human serum [25]. For example, the Caspian Sea has 132.44mM sodium and 3.04 mM potassium concentration [25] that is similar to the human serum (with 130-145mM sodium and 3.5-5.3 mM potassium concentration) [26] and thus from this aspect could be a good target for screening this enzyme. The main aim of this study is to screen an extensive dataset of publicly available genome assemblies and MAGs to define the distribution of enzyme methioninase across different bacterial and archaeal phyla. In addition, we screen 3 metagenomes sequenced from brackish waters of the Caspian Sea [25] and 44 metagenomes originating from deep groundwater samples also within the brackish range of salinity [27] to target enzymes more adapted to the brackish salinity range. Our results discuss the distribution of methioninase with an eco-evolutionary perspective, shedding light on the unprecedented potential contribution of this enzyme in promoting and sustaining cooperation within the highly oligotrophic and anaerobic deep groundwater communities. Our eco-evolutionary perspective highlights the target environments for screening and shows that culture-based screening approaches are potentially missing a considerable diversity of this enzyme due to the prevalence of this enzyme among anaerobic lineages and its presence in syntrophic lineages, a limitation that could be rectified by using in-silico screening approaches.

**Figure 1.**
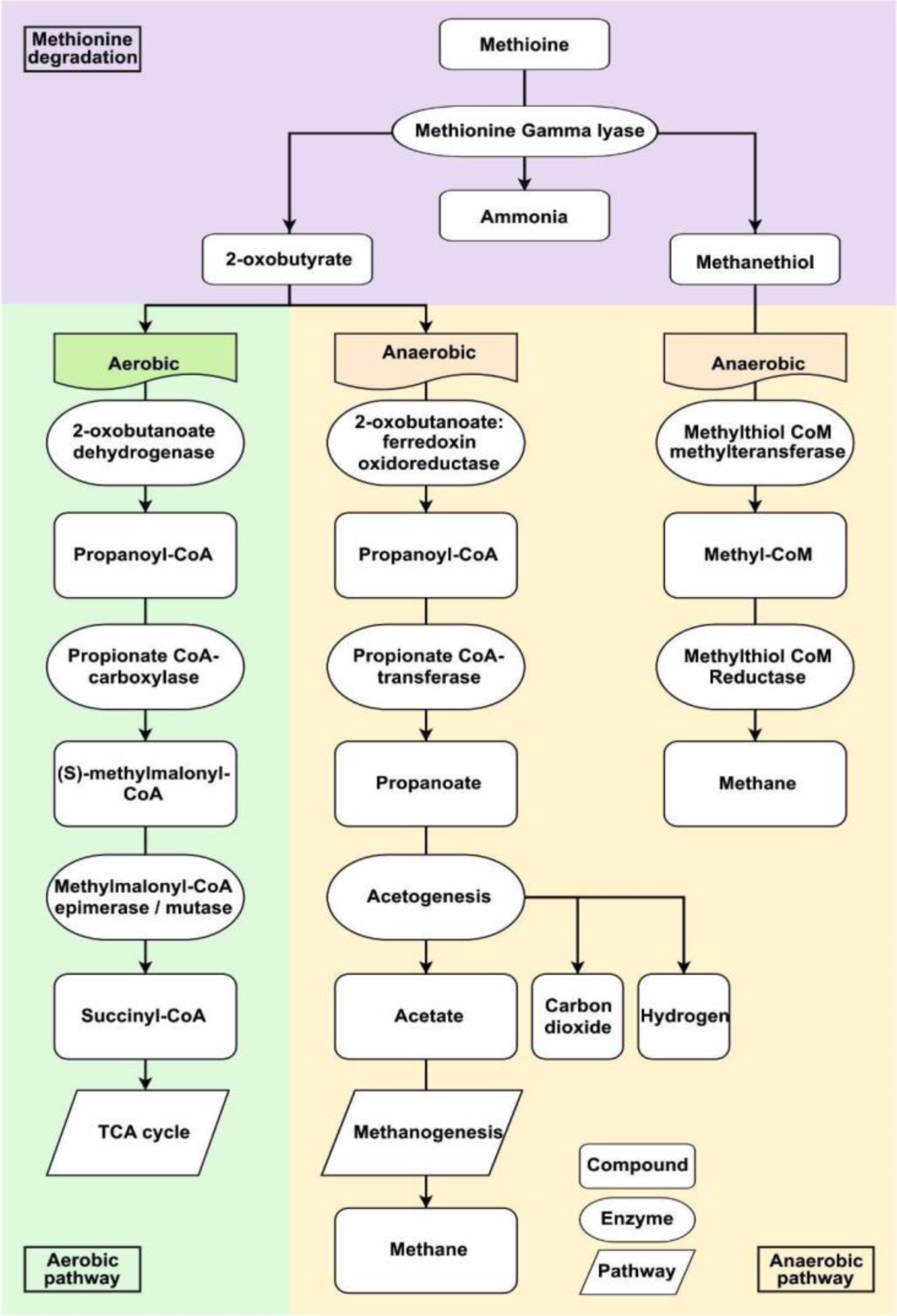
Catalytic pathways following methioninase activity on methionine. Catalytic activity of methioninase leads to the production of three products (i. e., ammonia, methanethiol, 2-oxobutyrate). The last two products could be further catabolized within cellular environment. In methanogenic archaea, methanethiol is utilized as substrate for methane production with methyl-CoM as an intermediate. However, two catalytic pathways are considered for 2-oxobutyrate based on aerobic or anaerobic metabolism of the host. In aerobic metabolism, 2-oxobutyrate is converted to propanoyl-CoA and then to methylmalonyl-CoA which is further processed to succinyl-CoA. This product is assumed to incorporate in the citric acid cycle. With anaerobic metabolism, after conversion of 2-oxobutyrate to propanoyl-CoA, propanoate is formed which is converted to acetate through acetogenesis process. Finally, acetate could be used also as a substrate for methane production.

## Results and discussion

### In-silico screening for methioninase

We performed an extensive in-silico screening of the enzyme methioninase in publicly available bacterial and archaeal genome assemblies and in a selected set of brackish metagenomes (sequenced from the Caspian Sea and Deep groundwaters of the Fennoscandian Shield). We used a methionine gamma-lyase (i. e., methioninase) Hidden Markov Model (HMM) from TIGRFAM (TIGR01328) as the search input to screen an extensive set of bacterial and archaeal genome assemblies (33,469 bacterial and archaeal genome assemblies) via the Annotree database (version 1.2.0, with taxonomy based on R95 GTDB release) [28]. We also used TIGR01328 to screen predicted open reading frames from the assembled depth profile metagenomes of the Caspian Sea [25], [29] as well as 1278 metageomes assembled genomes (MAGs) and 114 single-cell amplified genomes (SAGs) of the Fennoscandian Shield Genomic Database (FSGD) [27] for putative methioninases.

A total of 2,346 bacterial and 40 archaeal protein sequences were identified as putative methioninases via Annotree. All these annotations were further manually checked via the NCBI Conserved Domain Database (CDD) [30], [31], Blast KOALA [32] and HMMER [33] online servers. CDD annotated most sequences in aspartate aminoteransferase (AAT) superfamily (63%), that encompasses the enzyme methioninase. 25% of sequences targeted by TIGR01328 were annotated as cystathionine gamma-synthase, which is highly similar to methioninase [34], [35]. Few sequences (0.01%) were annotated as PLP dependent enzymes involved in cysteine and methionine metabolism (a broad annotation category also including both cystathionine gamma-synthase and methionine gamma-lyase). The remaining sequences (10%) were categorized as groups closely related to methionine gamma-lyase with the exception of only one sequence that was annotated as dinitrogenase enzyme (**Supplementary Table S1**). These mixed results from the CDD search hint at a high similarity between methioninase and other members of the AAT superfamily specifically cystathionine gamma-synthase; thus, necessitating a more precise approach for in-silico screening of methioninase. Annotations derived from BlastKOALA and HMMER also corroborated the CDD results.

The active site of enzyme methioninase contains two stretches of conserved amino acids critical for its specific activity. Cys116 of methioninase from *P.putida* is a major indicator of enzyme specificity. The mutation of Cys116 to histidine drastically reduces enzyme’s activity towards methionine. In addition, Tyr114 mutation in combination with Cys116 could completely eliminate the replacement reaction between methionine and 2-mercaptoethanol [36]. Inspecting methionine gamma-lyase sequences included in the seed alignment of TIGR01328, we could see that the stretch of Tyr, Gly and Cys (YGC) amino acids is conserved among them. In addition, Lys 240 and Asp241 (KD) are also key amino acids in the formation of methioninase active pocket [36]. Mutating these conserved residues mostly results in substantial decrease of the alpha/gamma elimination reaction [36]. Filtering out sequences missing YGC and adding the presence of KD amino acids as the second filter; from 2,346 bacterial entries and 40 archaeal entries, 1,753 and 18 protein sequences, respectively, were retained as putative methioninase. Despite being picked by the model and confirmed through the manual checking of the annotation, some sequences (22.4%) did not contain the conserved amino acids distinctive of methioninase.

The same in-silico search pipeline was applied to 2,775,993 predicted open reading frames (ORFs) of the assembled Caspian Sea metagenomes [25]. Filtering putative methioninase sequences missing YGC stretch left us with 7 entries and filtering those missing KD amino acids as the second step resulted in only 2 protein sequences (**Supplementary Table S1**). These two hits show a 100% protein sequence identity to each other. However, they originated from metagenomes sequenced from samples collected at two different depths of the Caspian Sea (40- and 150-meter depths).

Applying the same screening pipeline, a total of 72 methioninase hits were annotated and confirmed from 1278 MAGs and 114 SAGs reconstructed from deep groundwater metagenomic samples collected from the Fennoscandian shield which is a considerable leap from the single methioninase screened from the Caspian Sea metagenome (**Supplementary Table S1**). Activity of methioninase could lead to energy production in anaerobic metabolism [37] (**Figure 1**) and the anoxic environment of Fennoscandian shield deep groundwaters as compared to the fully oxic waters column of the Caspian Sea could provide methioninase-containing microbes with a competitive edge, potentially explaining the higher prevalence of this enzyme in the deep groundwater.

The screening pipeline for enzyme methioninase specifically highlights the importance of in-depth manual checks in order to confirm annotations retrieved from different models and databases. Despite applying threshold to the validity of hits detected by different models, in-silico annotations remain highly intricate specifically for closely related enzymes. Our results show that approximately less than 69.95% of annotations that were confirmed by multiple databases actually contain the amino acids in their active sites that are necessary for specificity and activity towards methionine.

### Distribution of Methioninase across domain Bacteria and Archaea

The majority of confirmed screened methioninase sequences belong to representatives of phyla Proteobacteria (50%), Firmicutes (29%), and Firmicutes_A (13%) (**Figure 2a and 2c**). The methioninase positive genomes in these phyla comprise 20%, 10%, and 5.47% of all representative genomes of the mentioned phyla within the screened genome assemblies, respectively (**Figure 2a and 2c**). The rest of confirmed screened methioninases (n=150) were detected in genomes belonging to 19 different phyla (i. e., Firmicutes_C, Firmicutes_E, Firmicutes_B, Firmicutes_C, Firmicutes_F, Firmicutes_D, Fusobacteriota, Patescibacteria, Bacteroidota, Acidobacteriota, Desulfobacterota, UBA10199, Synergistota, Riflebacteria, Spirochaetota, Eisenbacteria, AABM5-125-24, FEN-1099, DTU030, and Nitrospinota_B) (**Figure 2a and 2c**).

**Figure 2.**
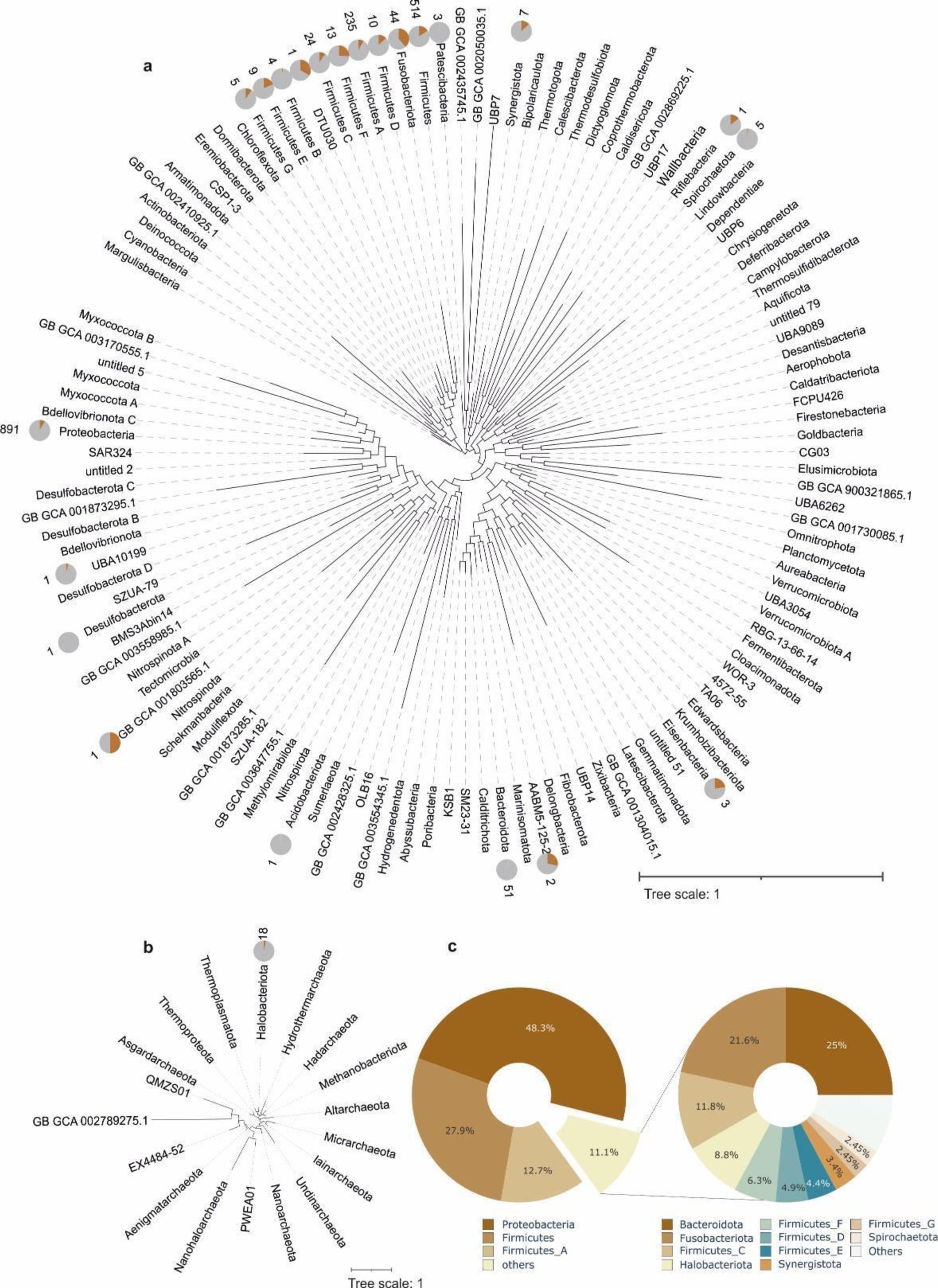
The phylogenetic distribution of screened methioninases. Phylogenetic distribution is shown across bacterial (a) and archaeal (a) phyla. Pie charts positioned in front of each phylum in panel a represent the number of genomes containing methioninase relative to all genomes within that phylum. The numbers presented in front of each pie chart indicate the count of genomes in which methioninase was detected. The large pie charts (c) depict the percentage of methioninase identified in each microbial phylum.

Interestingly, the 44 confirmed screened methioninases detected in representatives of the phylum Fusobacteriota (**Figure 2a**) highlight that close to half of available genomes affiliated to this phylum contain the methioninase enzyme. This relatively high prevalence of methioninase in representatives of this phylum could be related to their asaccharolytic metabolism relevant for their niche in the oral cavity [38]. Asaccharolytic microorganisms are forced to consume uncommon carbon sources such as amino acids [39]. Additionally, it is reported that the methanethiol produced via methionine degradation (**Figure 1**) in the bacterial population living in gingival sulcus, such as *Fusobacterium nucleatum*, could facilitate inflammation of gingival epithelium by increasing the permeability of these cells towards bacterial antigens, which makes methioninase not only a means for energy production, but also a virulence factor for these microorganisms [40].

At higher taxonomic resolution, relatively high prevalence of methioninases in each phyla are detected among representatives of classes *Halanaerobiales* in phylum *Firmicutes F (100%)*, *Clostridium* in phylum *Firmicutes A* (95.7%), *porphyromonas* in *Bacteriodota (59.4%)* and Negativicutes in phylum Firmicutes C (100%) (**Supplementary table S1**). A common metabolic feature among most representatives of these lineages is their anaerobic metabolism. Methioninase activity leads to the generation of three compounds: ammonia, methanethiol, and 2-oxobutyrate, with the latter two having the potential for being further catabolized (**Figure 1**). 2-oxobutyrate can be metabolized to propanoyl-CoA. While typically utilized for cellular component synthesis or energy production through pathways such as succinate-propionate [42], in anaerobic microorganisms engaged in symbiosis with methanogens, propanoyl-CoA can be converted to acetate [43], [44]. This acetate can then be transported to methanogenic microorganisms via an acetate transporter, where it is activated to acetyl-CoA. Acetyl-CoA undergoes dismutation to a carbonyl group, which is subsequently oxidized to CO_2_ and the methyl group is integrated into the methanogenic pathway. In aceticlastic methanogenesis, both membrane-bound methyl transferase and electron transport are crucial for energy conservation [45].

In archaea (**Figure 2b**), all of the confirmed screened methioninase sequences were detected in representatives of phylum Halobacteriota, class *Methanosarcina* (n=16) and class Syntropharchaeia (n=2).

Under anaerobic conditions, both methanethiol and 2-oxobutyrate can serve as substrates for methanogenesis (**Figure 1**), allowing for energy conservation throughout the process. Methanethiol is activated to methyl-CoA by methylthiol-CoM methyltransferase, resulting in the production of 3 methanes and one carbon dioxide from four methyl groups. The membrane-bound electron transport chain can utilize reduced ferredoxin and heterodisulfide, generated via the methanogenesis process, to establish a proton motive force for energy conservation [41]. The observed distribution of methioninase in methanogenic archaea is in agreement with our hypothesis of close connection of methioninase with methanogenesis capability [37], [46], [47]. Our results show that anaerobic metabolism and methanogenesis pathway are prevalent among genomes containing enzyme methioninase. In addition, the higher prevalence of methioninase in the deep groundwater samples reiterates that anoxic environments, that foster the proliferation of anaerobic microorganisms, could be a highly promising target to screen for novel methioninases.

### Functional validation for the Caspian Sea methioninase

Detecting only a single confirmed methioninase sequence among all predicted ORFs of the Caspian Sea depth profile metagenomes that are also oxic show a very low prevalence for this enzyme in these brackish waters. Capability to metabolize different amino acids shows a different distribution across different taxa. Amino acids such as glutamic acid and aspartic acid seem to be more commonly metabolized compared to methionine; specifically, since they could be easily integrated into the citric acid cycle [19]. Previous studies on screening of enzymes L-asparaginase and arginine deiminase (also with potential application for treating auxotrophic cancer cells) from metagenomic samples of the Caspian Sea, have resulted in relatively more hits for those enzymes (87 and 27 confirmed annotations for L-asparaginase [48] and arginine deiminase (results not yet published), respectively) as compared to the single hit recovered here for methioninase. An increasing prevalence along the depth profile of the Caspian Sea is detected for these enzymes (**Supplementary Figure S2**) which is in line with reports on proteolytic metabolism being more prevalent at deeper strata of aquatic environments [49]. For the case of methioninase where only one candidate was confirmed, it is also present at the deeper layers of 40 and 150 meters.

To validate the activity of this single enzyme screened from the oxic and brackish waters of the Caspian Sea, the coding sequence of the potential methioninase enzyme from the Caspian Sea metagenomes (3D protein structure in **Supplementary Figure S3**) was chemically synthesized. The codon optimized sequence was cloned in the pET26-b (+) in fusion with hexahistidine tag at the C-terminus and expressed in *E. coli* BL21(DE3). After optimizing the expression condition, maximum specific activity of crude extract was achieved by induction of mid-log phase transformed cells for five hours at 30°C using Luria-Bertani (LB) medium. Optimum Isopropyl b-d-1-thiogalactopyranoside (IPTG) concentration was 0.5mM. Recombinant methioninase was purified to homogeneity using nickel affinity chromatography. The enzyme sequence consists of 398 amino acids and the molecular mass of the monomeric enzyme based on the amino acid sequence was predicted to be 43kDa, which was consistent with protein migration on Sodium Dodecyl Sulfate PolyAcrylamide Gel Electrophoresis (SDS-PAGE) (**Supplementary Figure S4**). The protein sequence showed 57.58% identity to the methioninase from *P. putida* that has the same molecular weight. The molecular weight of bacterial methioninases varies usually in the range of 42.3 – 43.4 kDa according to SDS-PAGE studies [50].

The specific activity of the Caspian Sea methioninase was calculated to be 11.87 U/mg. Maximum specific activity of 520 U/mg has been reported for *P. putida*’s methioninase [24]. Specific activities of 21.9 and 182 U/mg are also stated for the methioninases purified from *Porphyromonas gingivalis* and *Arabidopsis thaliana*, respectively [40], [51]. These results further confirm the validity of our in-silico search via detecting in-vitro activity for the synthesized enzyme.

Moreover, the importance of identifying the suitable environment for each specific screening is emphasized. The low prevalence of methioninase among the Caspian Sea microbiome and the low activity of the single screened methioninase from the Caspian Sea metagenomes may be due to the prevailing oxic niches in this ecosystem. These oxic niches are not desirable for methioninase that is providing an edge to microbes growing under anaerobic conditions. Employing our approach for in-silico screening and functional validation via synthesizing confirmed sequences specifically for functions such as methioninase that are more prevalent among anaerobic microbes is highly promising. Via this approach, we not only bypass our severe limitations in culturing majority of the microbial community (great plate count anomaly) but specifically circumvent the need for screening under anaerobic condition that is additionally challenging.

### Phylogenetic relations of recovered methioninases

Examining the phylogenetic relations of protein sequences from all recovered putative methioninases by reconstructing an unrooted phylogeny provides an insight into the evolutionary relations of these enzymes. The confirmed methioninase sequence of the Caspian Sea branches together with other methioninases recovered from representatives of phylum Proteobacteria. Comparing this sequence against metagenome assembled genomes (MAGs) reconstructed from the Caspian Sea metagenomes [29] returns 100% sequence identity and full query coverage with a protein in the MAG casp40-mb.126 (GenBank assembly accession GCA_027618555.1). The taxonomic affiliation of this MAG based on GTDB is assigned to the phylum Proteobacteria (class *Gammaproteobacteria*, order *Pseudomonadales*, family *Porticoccaceae*, genus HTCC2207, and species HTCC2207 sp002382445) reiterating its close relation with other proteobacterial methioninases. All 5 publicly available genome assemblies in the genome taxonomy database (GTDB) affiliated to species HTCC2207 sp002382445 are reconstructed from brackish aquatic environment of the Baltic Sea (GCA_002382445.1, GCA_002430185.1, and GCA_937877205.1) and Caspian Sea (GCA_027618555.1 and GCA_027623455.1). Despite mainly oxic nature of these water columns; our in-vitro analyses confirm the activity for the screened methininase in this brackish lineage. Interestingly, mapping metagenomic reads on reconstructed Caspian Sea MAGs to assess the relative abundance of this MAG at different depths of the Caspian Sea; shows that the abundance of casp40-mb.126 in the 150 meters sample is close to double compared to the 40 meters sample (TPM of 1659 in 40 meters sample and 2987 in 150 meters metagenomes). Methioninases recovered from representatives of phylum Proteobacteria cluster in two main branches where one of them mainly contains representatives of order *Enterobacterales* and family *Alteromonadaceae*.

Phylogenetic reconstruction of recovered methioninases highlights some cases of potential horizontal transfer for the gene encoding this enzyme; where enzymes recovered from representatives of different phyla are clustering together. However, representatives of Proteobacteria and Firmicutes comprising the majority of recovered methioninase enzymes mainly form distinct clades.

All these putative methioninases included in this phylogeny contain key amino acids and belong to seven different groups based on CDD annotations (i.e., AAT_I superfamily, CGS_like, Cys_Met_Meta_PP, MetC, PRK06234, PRK06767, and PRK07503). Overlaying this information on the reconstructed phylogeny of the screened methioninases show that they are scattered across the tree (**Figure 3**). Interestingly, despite using TIGR01328 as search input and its presence among the NCBI CDD models, the best hit of the recovered sequences is to high-level annotations or taxa specific methioninase models such as PRK06767 (alignment representatives belong to Bacillaceae) detecting 39 hits affiliated to Firmicutes, and PRK07503 (alignment representatives belong to Proteobacteria) detecting 175 hits affiliated to Proteobacteria. Interestingly the in-silico screened and experimentally confirmed methioninase of the Caspian Sea is annotated at high level as AAT_I superfamily.

**Figure 3.**
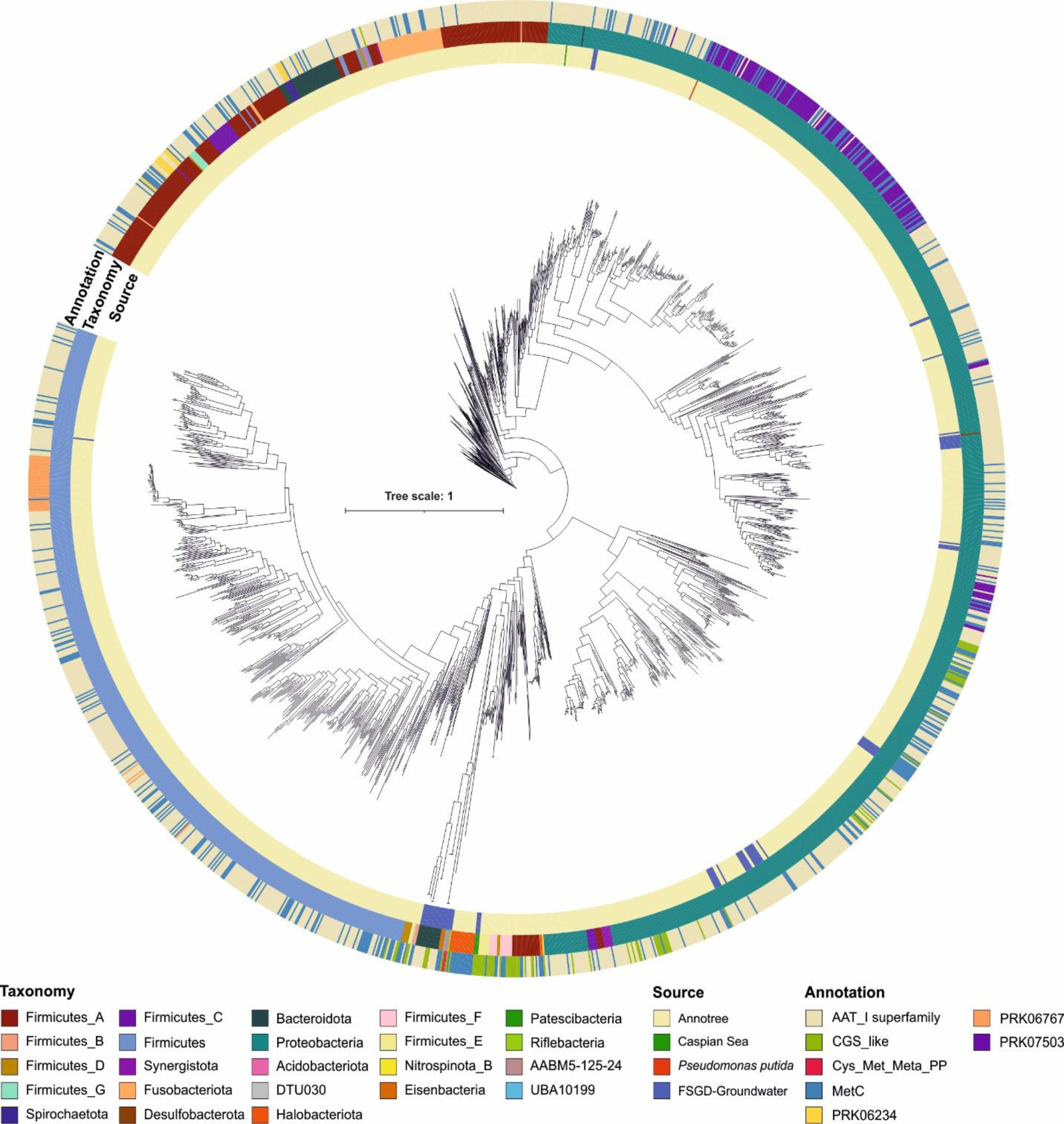
Phylogenetic relations of screened methioninases. Maximum likelihood phylogenetic relations of screened putative methioninases including the *Pseudomonas putida* methioninase sequence as the reference. The innermost color strip shows the source of the screened methioninase (from either Annotree, Caspian Sea, FSGD-groundwater, or Pseudomonas putida). The second color strip shows the taxonomic affiliation of the genomes containing each methioninase at phylum level. The outermost color strip shows the annotation derived from NCBI conserved domain database.

### The role of methioninase in metabolic syntrophy

An interesting observation from the groundwater MAGs/SAGs is the detection of 3 putative methioninase in three representatives of phylum Patescibacteria (two SAGs and one MAG affiliated to class Gracilibacteria, order Peregrinibacterales). The two SAGs and one MAG containing methioninase belong to the same species (average nucleotide identity ≥95%) and their assembly size is in the range of 1011935 to 1456149 with genome completeness in the range of 50.9 to 70.8 (details in the **Supplementary Table S2**) [27]. These three putative methioninase sequences are identical and have only 39.8% identity with the methioninase of *P. putida*. These three sequences were annotated as CGS_like via the NCBI CDD and K01761 (methionine-gamma-lyase [EC:4.4.1.11]) via BlastKOALA. Within their sequence, they contained the YGC and KD stretches of amino acids that are needed for the methioninase activity.

Representatives of phylum Patescibacteria are known for their small genome size as well as their fragmented metabolic capabilities [52]. It is due to their patchy metabolic potential that they are hypothesized to rely on a symbiotic lifestyle, most likely with other bacteria or archaea in the ecosystem. Inspecting metabolic capabilities of these three MAGs show that they do not encode genes involved in the methanogenesis pathway in order to utilize the products of methioninase activity. However, they seem to encode genes for this enzyme in the context of their otherwise fragmented metabolism. This potentially hints that methioninase could be playing a role in their metabolism in the highly oligotrophic deep groundwater ecosystem.

It is worth mentioning that Patescibacteria are present in a variety of ecosystems (e.g., deep lake strata, sediment environments, groundwater, etc.) and are not specifically underrepresented among the publicly available database of MAGs screened for methioninase via annotree (1,131 genome assemblies affiliated to Patescibacteria) and FSGD (319 genome assemblies affiliated to Patescibacteria). Apart from these 3 MAGs, only one other Patescibacteria MAG (GB_GCA_002329205.1) also contained a gene that was picked by the TIGR01328 model and was annotated as CGS_like via the NCBI CDD. However, this sequence was discarded since it did not contain the YGC and KD stretches of amino acids.

Annotating putative methioninases among the deep groundwater Patescibacteria underscores the potential role of this enzyme for symbiotic interactions among members of the microbial communities inhabiting this nutrient-poor environment. Presence of methioninase provides a potential advantage in syntrophic relationship of these Patescibacteria with other microbes capable of methanogenic metabolism that are among the main primary producers in these ecosystems [53]. There are studies showing that Patescibacteria are associated with methanogenic archaea [54]. Here we hypothesize that the encoded methioninase in these Patescibacteria affiliated MAGs could contribute in their syntrophic interactions where their methanogenic partners/hosts could potentially benefit from products of methionine degradation specifically the produced 2-oxobutyrate and methanethiol. These three methininases are branching just outside the branch containing all 16 methioninases annotated from representatives of class *Methanosarcina*, that are methane-producing archaea (**Figure 3**). However, detecting this protein in two single cell amplified genomes as well as a metagenome assembled genome confirms that this detection is not dues to a binning error. Interestingly among all 44 metagenomes sequenced from the deep Fennoscandian Shield [27] these three assemblies are only detected in one metagenome sequenced from the samples collected from borehole KA3385A [27]. This borehole contains groundwater from 448.35 meter below surface and has the salinity of 13.5 PPT. This high salinity hint at the long retention time of water in these depths (approximation at the scale of thousands of years) [27] (**Supplementary Table S2**). These deep and saline groundwater samples are among the most energy and nutrient limited ecosystems where methanogens could occupy a critical niche.

## Conclusions

By catalyzing the degradation of a sulfur containing amino acid, methioninase generates end products that put it at the interface of carbon, nitrogen, sulfur, and methane metabolism. Annotation of methioninases is challenged by low specificity of models that have high annotation ranks containing several similar enzymes. Moreover, some of these enzymes are reported to have different targets that is further complicating the *in-silico* search. By including multiple manual checks and presence of critical amino acids as pre requirements, we only detected and approved the annotation of 1845 methioninases from 35,338 screened genome assemblies. The low detection rate compared to other amino-acid degrading enzymes such as L-asparaginase prompted us to look into the metabolic capabilities of those encoding this gene. Prevalence of this enzyme among anaerobic microbes and its contribution to energy preservation under anaerobic metabolism highlights anoxic environments as an ideal target for screening novel methioninases. Our results highlight that niche aware screening is an important consideration for unleashing large-scale analyses. Additionally, it is via our refined annotation pipeline that we detect methioninase in representatives of Patescibacteria recovered from deep gruondwater. This finding highlights the unprecedented role of methioninase as a mean for syntrophic interactions of Patescibacteria with their host/partner.

## Methods

### Sequence assembly and annotation

Three publicly available metagenomic samples obtained from the brackish waters of the Caspian Sea at depths 15, 40 and 150 meters were used in this study [25]. The quality of sequencing reads was inspected with bbduk.sh [55] and MetaSPAdes [56] was used for assembly. Open reading frames (ORFs) were predicted using PRODIGIAL gene prediction tool [57]. A total of 586882, 1055141 and 1133215 ORFs were recognized in samples of 15, 40 and 150 meters, respectively. Metagenome assembled genomes (MAGs) and single cell assembled genomes (SAGs) reconstructed from the deep groundwater samples of the Fennoscandian shield were retrieved from the Fennoscandian Shield Genomic Database (FSGD) [27]. For each MAG and SAG open reading frames were predicted using PRODIGIAL gene prediction tool [57].

### Methioninase screening

Methioninase HMM model was acquired from the TIGRfam database with the ID of TIGR01328 [58]. This HMM model was used to search annotated genomes of the Annotree via its web service (version 1.2.0 using R95 GTDB release) [28]. A total of 2346 bacterial and 40 archaeal putative methioninases from respectively, 45 and 3 phyla were recovered from the Annotree database. These sequences were manually validated by annotation against Conserved Domain Database (CDD) [30], [31], HMMER sequence database (hmmscan analysis) [33]and BlastKOALA databases [32]. The hmmscan annotated all of the presented sequences as “Cys/Met metabolism PLP-dependent enzyme” while BlastKOALA annotated 2363 sequences as “methionine-gamma-lyase [EC:4.4.1.11]” and 23 as “CTH; cystathionine gamma-lyase [EC:4.4.1.1]”. However, CDD database annotated more than half of these sequences as a member of “AAT I superfamily” and the rest as a part of cd00614, CGS_like, cl18945, COG0626, Cys_Met_Meta_PP, MetC, pfam01053, PRK06234, PRK06767 or PRK07503 groups. Not all of these groups link to methioninase but they are close relatives or at a higher order of annotation hierarchy. Only one sequence, PMZT01000246.1, was annotated as MetC as well as “NifX_NifB”, which is a family of iron-molybdenum cluster-binding proteins.

For screening the Caspian Sea metagenomic sample, with hmmscan of the HMMER analysis package and the same methioninase HMM model from TIGRfam database, putative methioninase sequences were screened from metagenome derived ORFs. The threshold was set at 150 to capture distant enzymes with activity similar to methioninase. Annotation of these sequences with BlastKOALA resulted in 7 protein families, methionine-gamma-lyase, cystathionine gamma-lyase, cysteine-S-conjugate beta-lyase, O-acetylhomoserine (thiol)-lyase, O-succinylhomoserine sulfhydrylase, cystathionine gamma-synthase and O-acetylhomoserine/O-acetylserine sulfhydrylase. Among these, 144 sequences annotated as methionine-gamma-lyase were selected for further analysis. CDD annotation divided these 144 sequences into 4 groups, 97 sequences were annotated as a member of AAT_I superfamily (cl18945), 22 as Cystathionine gamma-synthase like protein (cd00614), 25 as Cystathionine beta-lyase/cystathionine gamma-synthase (COG0626) and the last 2 sequences were annotated as MetC superfamiliy (cl43216).

According to Fukumoto and colleague’s findings [36] tyrosine 114, glycine 115 and cystine 116 (YGC) in combination with lysine 240 and asparagine 241 (KD) are the key amino acids involved in methioninase function. Backed by this knowledge, these residues were used as a guide to find sequences that are more likely to show a great activity towards methionine. Seven sequences contained YGC residues and only 2 sequences, which are identical to each other, contained YGC and KD residues in the proper locations. The same manual filtration based on methioninase conserved amino acid stretches was performed on the sequences from annotree, which resulted in 1771 methioninases.

For the Fennoscandian Shield Genomic Database MAGs and SAGs, Utilizing PRODIGIAL, all their ORFs were predicted and their sequences were used as an input for HMMscan with the methioninase HMM model from the TIGRfam database. With this HMM model, 5890 sequences were initially annotated as methionine gamma-lyase. However, BlastKOALA only confirmed 246 of these sequences as methionine gamma-lyase. Same as before, CDD annotations showed more diversity and 106 sequence were annotated as a member of AAT_I superfamily (cl18945), 66 sequences were annotated as cystathionine gamma-synthase (cd00614), 21 sequences as enzymes involved in cysteine and methionine metabolism (pfam01053), 16 sequences as a member of cystathionine gamma-synthase (PRK07049) and finally 37 sequences as Cystathionine beta-lyase/cystathionine gamma-synthase (COG0626). Following the same pipeline as before sequences containing both YGC and KD residues at the proper location were selected as true methioninases, resulting in 72 sequences.

### Distribution of methioninases across different phyla

Distribution of all methioninase sequences retrieved from Annotree, and those screened from the Caspian Sea and groundwater samples of the Fennoscandian Shield metagenomes was represented across different phyla and classes. The base of the phylogenetic trees was obtained from the Annotree database as a newick file. To show the relative abundance of methioninase, the number of genomes containing methioninase and the total number of genomes represented in each bacterial phyla (class level in the archaea tree) was represented as a pie chart in front of each taxon using iTOL web server (**Figure 2**) [59].

### Reconstructing phylogenetic relations of recovered methioninases

All of the confirmed methioninase sequences retrieved from Annotree, screened from the Caspian Sea and the Fennoscandian Shield ground water metagenomes were aligned using Kalign (v 3.0) using 10.0 gap opening penalty and 3.0 for gap extension penalty [60]. After this, aligned protein sequences were used for phylogeny reconstruction using FastTree [61].

### Gene cloning and enzyme expression

The single confirmed methioninse sequence screened from the Caspian Sea metagenomes was used for gene synthesis. The codon optimized synthetic methioninase sequence was cloned in the NdeI-XhoI restriction sites of pET26-b(+)in frame with the C-terminal His-tag, and transformed into *E. coli* BL21(DE3). The right transformant based on sequencing results, was inoculated into the LB medium supplemented with 50 µg/ml kanamycin at 37℃ until reaching optical density (OD) of 0.6; an overnight grown pre-culture was used for this purpose. Different concentrations of IPTG (0.1, 0.3, 0.5, 0.8 and 1 mM), induction duration (3, 5 and 7 h and overnight), induction temperature (20, 25 and 30 °C) as well as the culture media (Luria Bertani (LB) and M9) were tested to find the optimal expression condition. Finally, protein expression was induced by addition of 0.5 mM IPTG at 30°C for **5** h.

In the following step; cells were spun down (9000 g, 10 min), concentrated in 5 ml of ice-cold lysis buffer (50 mM potassium phosphate buffer, pH 7.5, NaCl 300 mM and Urea 2M) and lysed by sonication (recurrent periods of 10s pulsing and 5s resting for 10 minutes, SYCLON Ultra Sonic Cell Sonicator SKL950-IIDN). Lysate was centrifuged (9000g, 30 min, 4°C) and supernatant was loaded on Ni-NTA agarose column pre-equilibrated with lysis buffer. Elution of bound proteins was achieved by the buffer containing 250 mM of imidazole (pH 7.2). Purified protein fractions were dialyzed overnight against 50 mM phosphate buffer (pH 7.2, 4°C) to remove imidazole. All protein purification steps were carried out at 4°C.

SDS-PAGE (12% w/v acrylamide) was used to confirm the purity of the enzyme. Gels were stained by Coomassie Brilliant Blue R-250 (Bio-Rad, USA). Protein concentration was estimated by Bradford method and using bovine serum albumin as standard [62].

### Enzyme activity assessment

To quantify the amount of ammonia generated by the methioninase reaction, Nessler’s reagent was employed [63]. A standard curve was generated over a range of ammonium chloride concentrations (0.01 mM to 0.4 mM). Purified enzyme was added to 450 µL potassium phosphate buffer solutions (50 mM, pH 7.2) containing 200 mM of L-methionine and 0.01 mM pyridoxal 5-phosphate as methioninase cofactor. The reaction mix was incubated at 37℃ for 30 minutes prior to addition of 50 µL of Nessler’s reagent. Colorimetric assay was used to measure the amount of ammonia production at 420 nm. One unit (U) of enzyme’s catalytic activity was defined as the amount of enzyme which converts one micromole of L-methionine to ammonia per min at the condition of assay. The specific activity was expressed as U/mg protein. All assays were performed in triplicate, the mean and standard deviation were presented for each measurement.

## Data availability

The Caspian Sea metagenomes used for this study have been deposited to GenBank by Mehrshad et al. [25] and are accessible via the Bioproject PRJNA279271. All reconstructed MAGs from these metagenomes are also deposited to GenBank and are accessible under the accession number Bioproject PRJNA279271. Fennoscandian Shield genomic database (FSGD) MAGs and SAGs have been deposited to GenBank by Mehrshad et al. [27] and are accessible under the accession number Bioproject PRJNA627556. The MAGs and SAGs of FSGD are publicly available in figshare under the project “Fennoscandian Shield genomic database (FSGD)” with the identifier https://doi.org/10.6084/m9.figshare.12170313.

Alignment used for phylogeny reconstruction and the tree file as well as putative methioninase sequences are accompanying this manuscript as **Supplementary Data S1**.

## Acknowledgement.

The computational analysis was performed at the Center for High-Performance Computing, School of Mathematics, Statistics, and Computer Science, University of Tehran.

## Author contribution

M.M. and S.A. designed the study. E.KH. performed the bioinformatics analysis with supervision from M.M.. E.KH. and Z.V. performed the experimental analyses with supervision from S.A.. E.KH. and M.M. drafted the manuscript. All authors analyzed and interpreted the data and approved the manuscript.

## Supplementary

**Supplementary Figure S1.**
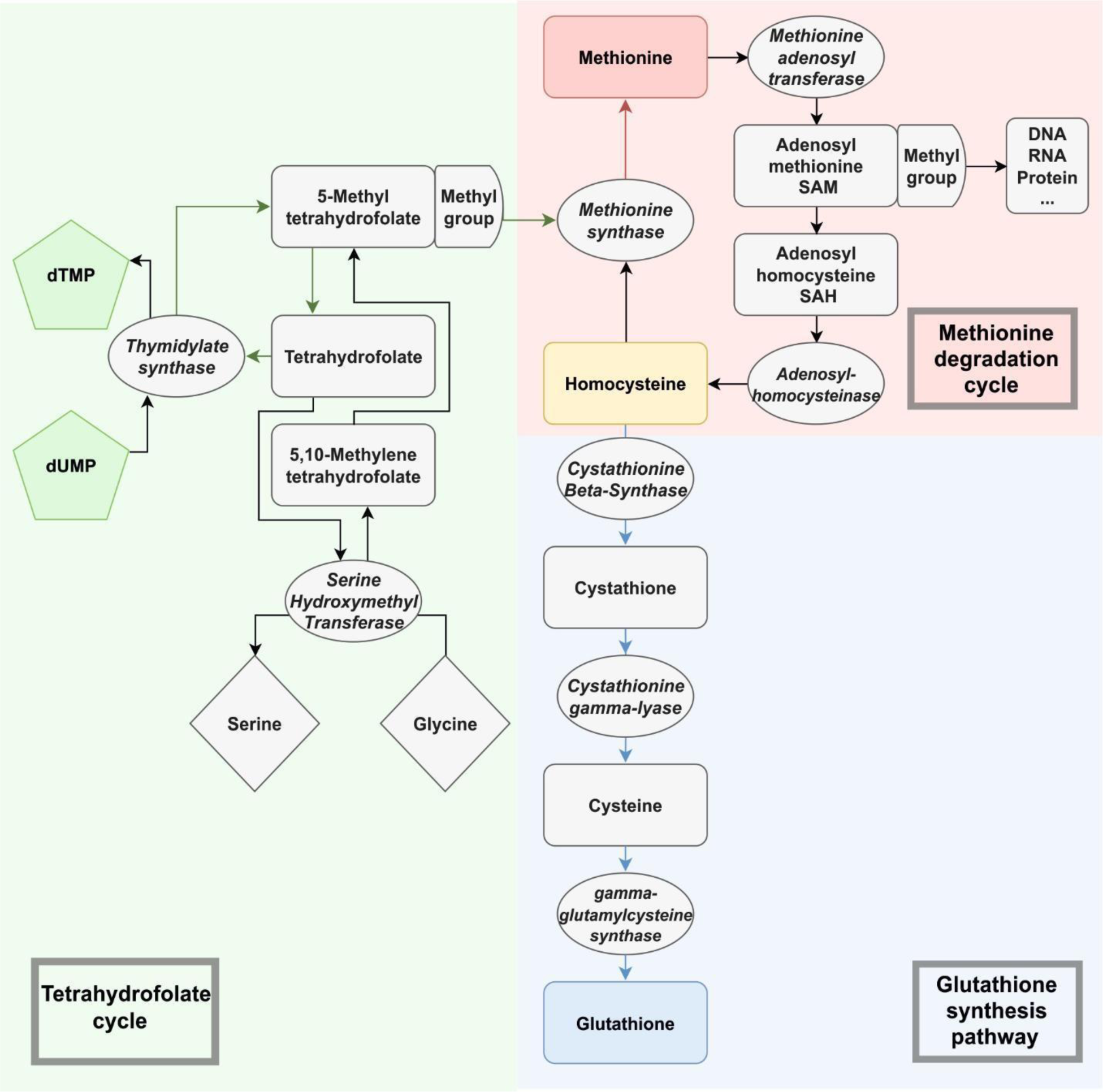
Dysregulated Pathways Influencing Methionine Production in Cancer Cells. In cancer cells, methionine production is compromised through three distinct mechanisms. Firstly, a significant portion of homocysteine is diverted towards the synthesis of glutathione, crucial for maintaining the cell’s redox homeostasis. Secondly, the production of 5,10-methylenetetrahydrofolate and 10-formyltetrahydrofolate, both essential intermediates in nucleotide biosynthesis, exhibits an inversely proportional relationship with methionine production. Lastly, S-adenosylmethionine (SAM), a pivotal product within the methionine cycle, is predominantly utilized as a methyl donor for DNA, RNA, and protein methylation in cancer cells. Together, these three pathways contribute to the development of methionine auxotrophy in cancer cells.

**Supplementary Figure S2.**
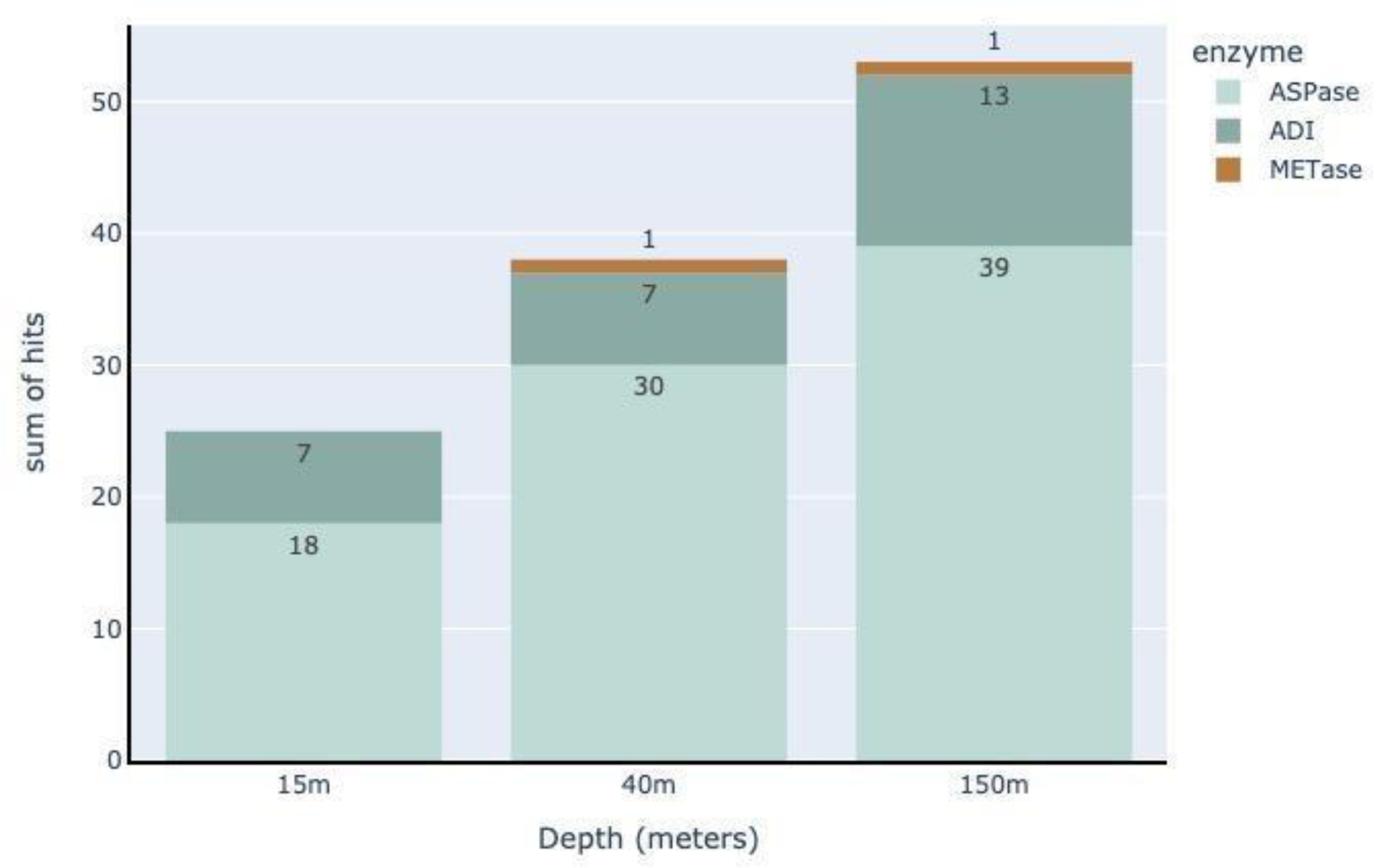
Distribution of asparaginase, arginine deiminase and methioininase in metagenomic samples collected from different depths of the Caspian Sea brackish water.

**Supplementary Figure S3.**
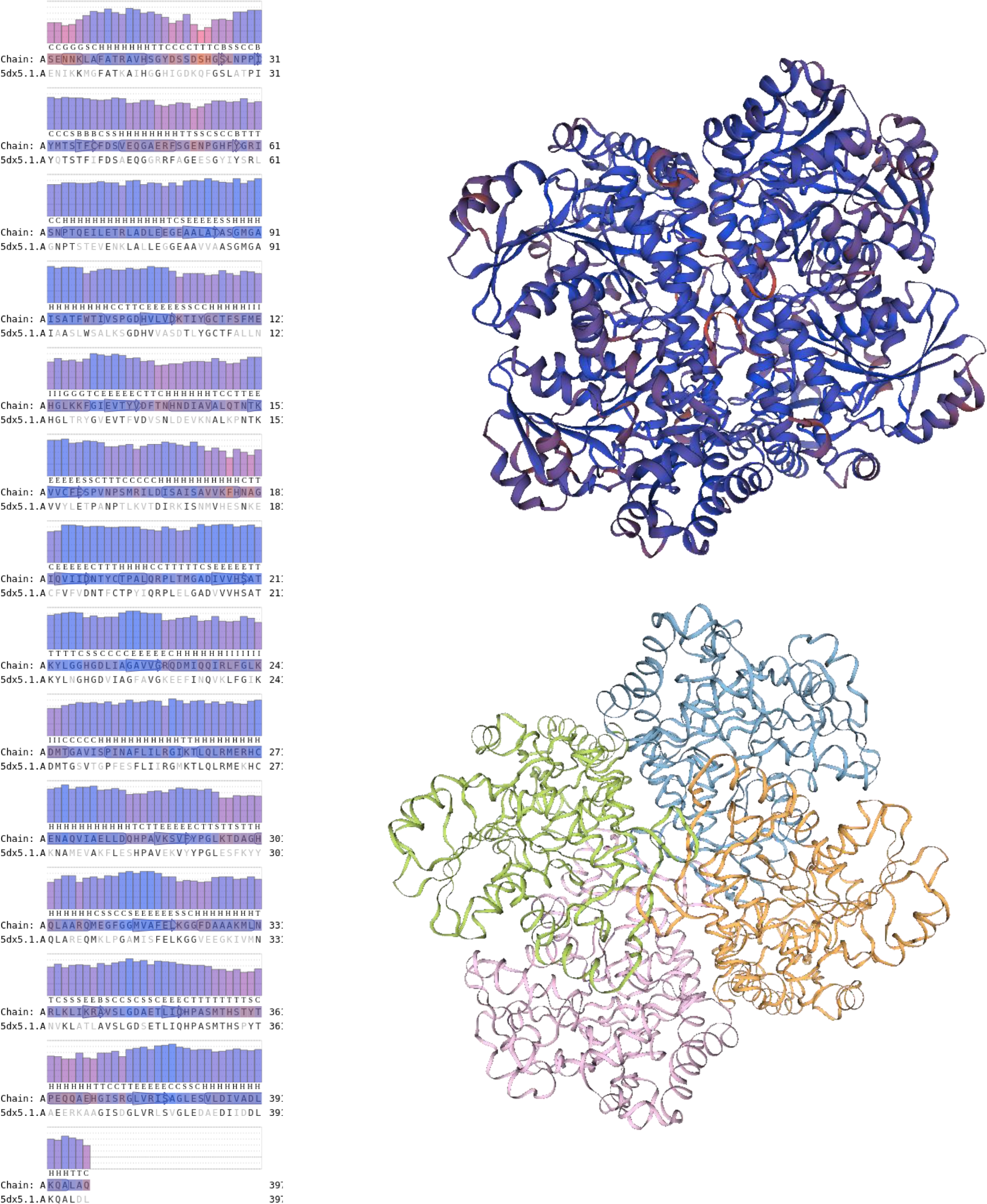
predicted 3D structure of the methioninase screened from Caspian Sea metagenome. The alignment (left) depicts the high similarity of the screened methioinase protein sequence with the Clostridium sporogenes (strain ATCC 15579) methioninase (5dx5).

**Supplementary Figure S4.**
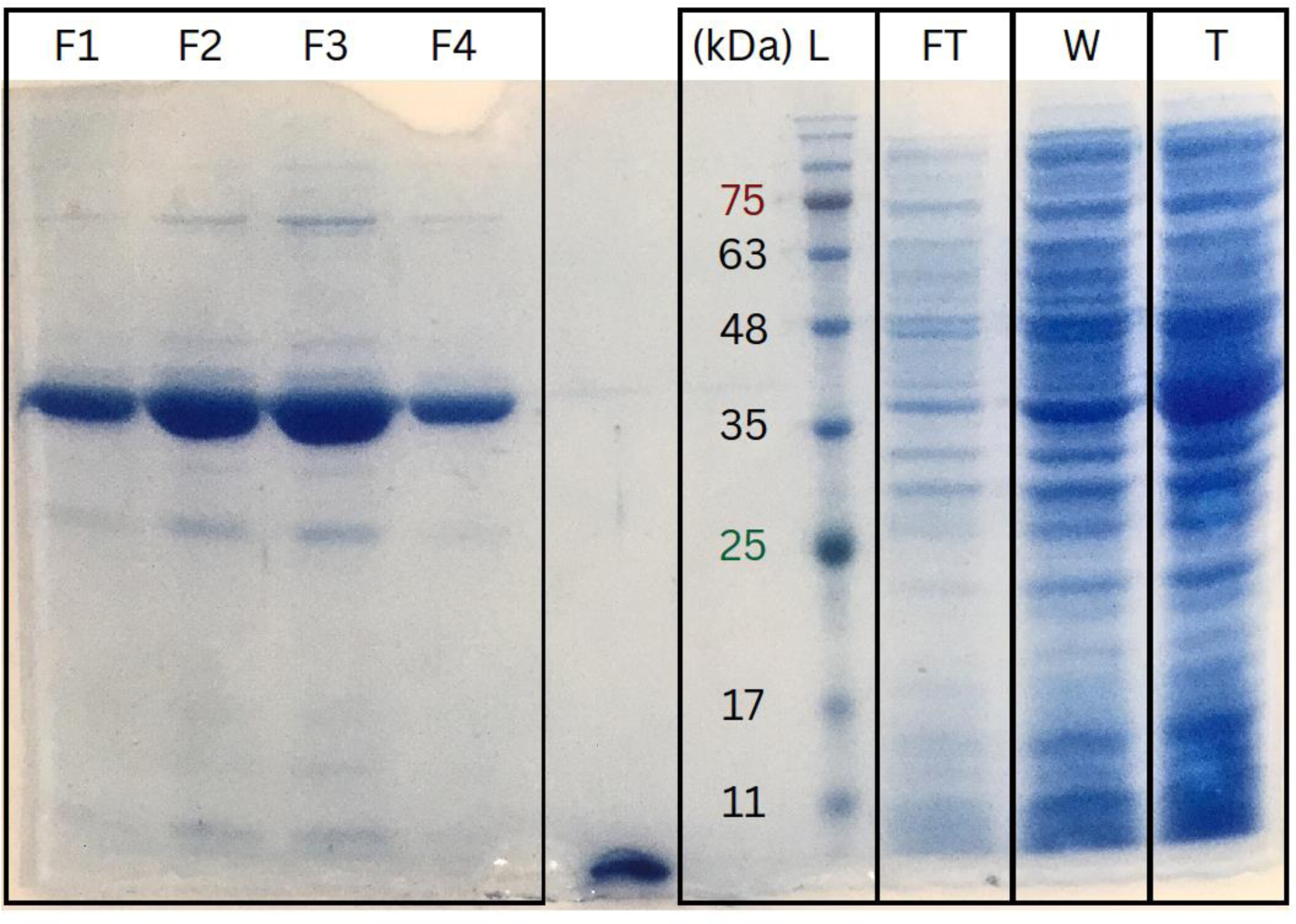
SDS-PAGE gel depicting the purification process of methioninase utilizing a Ni-NTA agarose column. From right to left, Total protein cell lysate (T), presenting a mixture of soluble proteins prior purification. Washed proteins (W), illustrating the removal of non-specifically bound proteins. Flowthrough (FT), indicating proteins that did not interact with the column. Protein ladder (L). Fractions of purified methioninase (F1, F2, F3 and F4), presenting the enriched methioninase. The patterns observed in this SDS-PAGE suggests a successful isolation and enrichment of methionianse.

